# Decline in Environmental Quality and Spatial Dynamics of New City Area Development Metropolitan Mamminasata, South Sulawesi, Indonesia

**DOI:** 10.1101/2022.01.28.478145

**Authors:** Batara Surya, Agus Salim, Syahrul Sariman, Hernita Hernita, Haeruddin Saleh, Seri Suriani, Nasrullah Nasrullah, Emil Salim Rasyidi

## Abstract

The expansion of urban areas towards the development of new urban areas has an impact on changes in land use, socio-economic dynamics and a decrease in environmental quality. This study aims to analyze land use change as a determinant of environmental degradation and the spatial dynamics of metropolitan urban areas, the effect of land use change, housing development, infrastructure development, and land reclamation on the environmental degradation of the new city area, and control strategies environmental pollution and sustainable development of the new city area. This study uses a quantitative-qualitative concurrent triangulation design approach. Elaboration of data through observation, in-depth interviews, surveys, and documentation is used to describe the socio-economic community, and the decline in the environmental quality of new city area development in relation to the spatial dynamics of metropolitan urban areas. The results of the study show that the intensity of land use change coupled with an increase in socio-economic activities in the development of new city areas is positively associated with a decrease in environmental quality and segregation towards the spatial dynamics of metropolitan urban areas. Changes in land use, housing development, infrastructure development, and land reclamation simultaneously affect environmental quality degradation with a coefficient of determination of 64.96%. This study recommends strategies for controlling environmental pollution and sustainable development of new city areas for the needs of formulating urban development policies for Mamminasata Metropolitan South Sulawesi, Indonesia.

## Introduction

Urbanization in the dynamics of the development of large and metropolitan cities in addition to having an impact on increasing population and pressure on space, also affects the availability of land and urban spatial dynamics. Urbanization is a driving force for spatial diffusion, increasing space requirements, and ecosystem complexity towards changes in the metropolitan urban structure [1–3]. The expansion of urban areas through the development of new urban areas has an impact on the conversion of productive agricultural land and shifts in the socio-economic status of the community. The dynamics of a metropolitan city that develops into suburban areas have an impact on the socio-economic conditions of the local community who originally occupied the land in the location of the new city area [4,5]. The dominant allocation of urban space utilization in fulfilling housing and settlement needs is positively associated with changes in land cover, spatial patterns, and the structure of urban space metropolitan. Thus, it is necessary to control the use of space towards a balance of physical, economic and environmental development [6,7]. Thus, the expansion of the city is a trigger for the movement of people from the city center to the suburbs. Suburbanization is a factor that affects the expansion of urban areas to suburban areas causing changes in spatial patterns due to increased development activities [8,9]. Thus, it is necessary to regulate the use of space and the allocation of landscape resources in a reasonable manner in order to achieve environmental balance towards sustainable development [10,11].

The orientation of urban development in Indonesia is dominantly developed towards new growth through urban expansion to suburban areas. This means that urban growth is influenced by three main factors, namely the process of capitalization, regional expansion or reclassification, and migration from villages to cities [12]. New cities in Indonesia were developed to meet the need for housing and socio-economic activities due to the increasing population. The implications of this development have an impact on the land cover index and changes in typology and morphology from rural areas to urban areas. This means that effective control over productive resources, especially land, is needed for sustainable development [13]. Thus, policy support is needed to balance development incentives and restrictive measures to ensure the sustainability of urban development [14,15]. Furthermore, the Mamminasata Metropolitan urban integration system contributes positively to population migration, circular migration, transportation generation and attraction and air quality pollution originating from motor vehicle exhaust gases. Urban transportation through the use of motorized vehicles is a major source of air pollution that requires treatment to reduce health risks for the community [16,17]. The development of new urban areas in suburban areas in addition to having an impact on the complexity of the ecosystem, also has an impact on the potential threat of flooding due to land reclamation carried out by housing developers and the community. The effects of different land use scenarios on subsequent changes in ecosystem services have major implications for sustainable land management [18]. This means that ecological security is important to maintain ecosystem functions and provide ecosystem services for human well-being [19]. Furthermore, the decline in environmental quality due to the intensity of the physical development of new urban areas is positively associated with the use of deep groundwater and the loss of biodiversity. Thus, the main challenges in the development of new urban areas are reduced hydrological functions, environmental pollution and excessive use of water resources [20,21].

The new city area of Moncongloe–Pattalassang was originally a productive agricultural area and a production forest area designed to support the growth of the Maminasata Metropolitan. The land used for development is 4,300 ha. The functions of urban activities developed include: (1) Housing and settlements occupying an area of 3,100 ha; (2) Commercial and service areas occupy an area of 50 ha; (3) Infrastructure development utilizes an area of 35 ha; and (4) Preparation of development land area of 800 ha. The allocation of built space in addition to having an impact on spatial dynamics also affects the socio-economic conditions of the suburban area community. Thus, the urgency of this research is focused on three important things, namely: (1) The effect of land conversion in the new city area on environmental quality degradation; (2) The complexity of spatial use and integration of urban systems has an impact on increasing traffic volume and population mobility based on the pattern of origin and destination of travel; and (3) Area expansion for the development of new cities has an impact on the utilization of water catchment areas and reduction of land cover. This study offers management of urban area development based on environmental pollution control and improvement of environmental quality and carrying capacity towards the sustainability of metropolitan suburban areas. Furthermore, environmental, economic, and social sustainability are important aspects needed in policy formulation and implementation of new urban area development as a unified metropolitan urban system. The concept of new city area development was created to anticipate excessive urbanization through the preparation of housing and settlements in suburban areas, distribution of economic activity services and reducing the high population density in the main city. The new city was created to overcome the problems of the main city in terms of fulfilling housing needs and socio-economic activities which were located in suburban areas [22,23]. Furthermore, this study examines the conceptual framework of the impact of land use change in the new city area and the integration of the Mamminasata Metropolitan urban system on environmental degradation. The conceptual framework for this study is presented in Figure 1 below.

**Fig.1.**
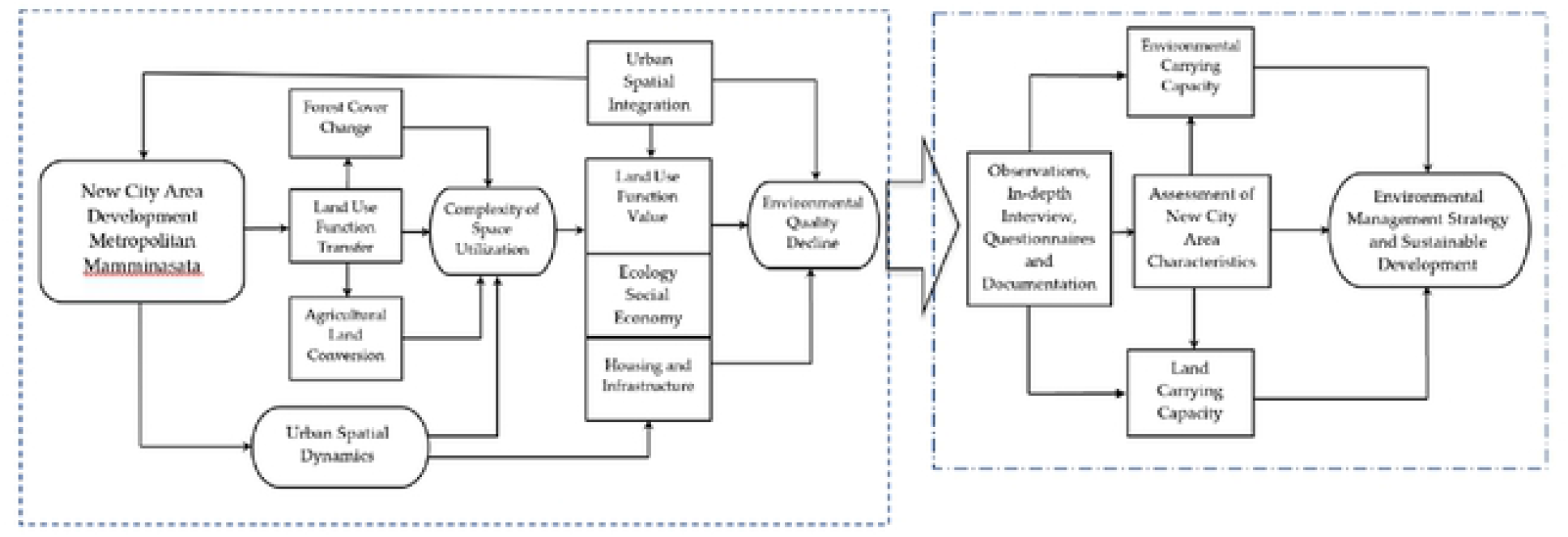
Conceptual framework of environmental degradation and spatial dynamics of new city areas. Source: Author’s elaboration

## Material and Method

This study uses a quantitative-qualitative concurrent triangulation design approach. Qualitative data obtained through in-depth interviews and documentation. Quantitative data were obtained through observation and surveys using a questionnaire instrument. Furthermore, the validation of the data in this study was carried out through a triangulation process. That is, this study uses data simultaneously in relation to the validity of the data obtained in the field. Thus, triangulation in the study was carried out by means, namely (i) triangulation of data sources, (ii) triangulation of theory, and (iii) extension of observation time.

### Study Area

This research was carried out from March to October 2021. The selection of the study location was based on the following considerations: (1) The development characteristics of the new city area of Moncongloe– Pattalassang contribute to the spatial dynamics and urban transportation system of the Mamminasata Metropolitan; (2) The allocation of developed space affects the characteristics of the surrounding rural areas; and (3) the development activities that are developed have an impact on the socio-economic changes of the community and the degradation of environmental quality. The spatial utilization of the new city area is presented in Table 1 below.

**Table 1.** Space utilization of the new city area of Moncongloe–Pattalassang in 2021

| Number | Land Use | Land Area (ha) | % |
| --- | --- | --- | --- |
| 1 | Mixed garden | 32.1 | 1.4 |
| 2 | Empty land | 194.5 | 9.3 |
| 3 | Production forest | 290.3 | 13.9 |
| 4 | Catchment area | 49.1 | 2.4 |
| 5 | Ricefield | 133.8 | 6.4 |
| 6 | Pond | 15.1 | 0.7 |
| 9 | Services | 12.7 | 0.6 |
| 10 | Medical facility | 2.1 | 0.1 |
| 11 | Education facility | 5.3 | 0.3 |
| 12 | Shopping center | 24.1 | 1.2 |
| 13 | Facilities of worship | 2.0 | 0.1 |
| 14 | Office | 1.7 | 0.1 |
| 15 | Settlement | 1,325.2 | 63.5 |

**Sumber:** Reference [22].

Table 1 shows that the dominant spatial uses in the new city area of Moncongloe-Pattalassang are settlements, production forests, vacant land for development, and agricultural areas (rice fields). Furthermore, activities that utilize the smallest land area are offices, religious facilities, and health facilities.

### Method of collecting data

#### Observation

Data collected through observations include: (1) Land reclamation and land preparation for new city areas; (2) Housing development and socio-economic activities that are built; (3) The structure and spatial pattern of the new city area; (4) Ecosystem conditions and characteristics; (5) Transportation system; (6) Environmental carrying capacity; and (7) Sources of environmental pollution. Furthermore, the instruments used in carrying out the observations are (i) field notes, (ii) periodic notes and (iii) check lists (check list data).

#### In-depth interviews

Data collected through in-depth interviews, among others: (1) Mechanisms and procedures for implementing the development of new urban areas; (2) land status and value; (3) The pattern of social and economic activities of the community; and (4) the social structure of society. The tools used in in-depth interviews, namely tape recorders, cameras, and interview guides equipped with freelance notes, checklists, and score scales.

#### Questionnaire

The data collected through the questionnaire include: (1) Changes in land use, measured by indicators, namely the area of built up land, socio-economic activities, space requirements, and land availability; (2) Housing development, measured by indicators, namely the area of land used, the type of housing developed, the basic building coefficient and building floor coefficient, land and building ownership; (3) Infrastructure development, measured by indicators, namely road infrastructure, drainage network, clean water pipe network, electricity network, and waste water treatment facilities; (4) Land reclamation, measured by indicators, namely land elevation, use threshold, reclamation procedures, and land carrying capacity; and (5) Environmental damage, measured by indicators, namely environmental quality index, source of pollution, potential for flooding, space capacity, and ecosystem stability. Measurement of the questionnaire data using the litkers scale, which is a value of 5 for the very supportive category, a value of 4 supportive, a value of 3 being enough supportive, a value of 2 being less supportive, and a value of 1 not supportive. The sample in this study was determined by purposive sampling, which was determined by the researcher with certain criteria. The actors who filled out the questionnaire included: (1) housing developers, (2) local government, (3) people living in the new city area, and (4) the community around the new city area who received the impact of the development. Furthermore, sampling refers to Cohran, [24]. The formulation used in the determination of the sample is as follows:

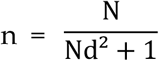

Where n is the number of samples, N is the number of population and d is the error rate (0.5) or 5% of the 95% confidence level. The number of samples set in this study is 250 respondents.

#### Documentation

The documentation used in this study includes: (1) Mamminasata Metropolitan Urban Spatial Plan Documents obtained from the Office of Human Settlements and Spatial Planning of South Sulawesi Province; (2) Documents for new city area infrastructure development are obtained through the Public Works Department of Highways of South Sulawesi Province; (3) Documents for the development of the new city area of Moncongloe–Pattalassang were obtained from the developers and the District Office.

#### Qualitative Data Analysis

Qualitative data analysis was carried out during field observations and at the time of separating information into categories. The stages of qualitative analysis carried out include: (1) Domain analysis is carried out by observing social situations, namely place, actor and activity (PAA). The aim is to determine the research domain, namely land use change works as a determinant of environmental degradation and urban spatial dynamics of the Mamminasata Metropolitan; (2) Taxonomic analysis based on the chosen domain to be described in more detail to understand the internal structure and developing social dynamics; (3) The compenesial analysis is carried out by contrasting the differences that arise in the dynamics of the development of the new city area of Moncongloe–Pattalassang; and (4) Cultural theme analysis is carried out by integrating across domains based on the focus and research objectives to be achieved. Stages of analysis carried out to support the results of this study, namely the measurement of the environmental quality index of the new city area of Moncongloe–Pattalassang.

#### Environmental Carrying Capacity Analysis

The carrying capacity of the environment is assessed based on the socio-economic dynamics of the community and the development activities developed. Parameters analyzed, among others: (i) carrying capacity of settlements, (ii) developer threshold, (iii) carrying capacity of land, (iv) carrying capacity of water resources, (v) carrying capacity of space and protected functions, (vi) index green open space, (vii) the carrying capacity of the environment, and (viii) the carrying capacity of ecosystem services. Furthermore, carrying capacity is defined as the ability and availability of land for the provision of residential land, infrastructure and other activities to accommodate the number of residents to live properly. The analysis of the carrying capacity of settlements uses the following formulation:

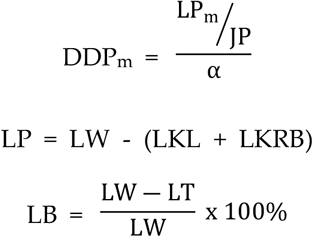

Where DDP_m_ is the carrying capacity of settlements, JP is the number of residents, *α* is the coefficient of the area of space requirements (m^2^ / capita), and LP_m_ is the area of land suitable for settlement (m^2^). LW is the area, LKL is the area of protected areas and LKRB is the area of disaster-prone areas. LB is building coverage (%), LW is the area of the new city area, and LT is open space. Limits for the feasibility of carrying capacity of land for settlements, namely (i) DDP>1 means able to accommodate residents to live, DDP=1, means there is a balance between the population living with the area, DDP,1 means unable to accommodate residents to live. Note: if the value of LB = 0% means that the land has not been used for buildings, otherwise if the value of LB = 60%, then the use of land for buildings reaches 60% of the total area. Furthermore, the threshold for settlement development uses the following formulation:

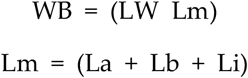

Where WB is an area that can be developed (ha), LW is the area of a new city area (ha), Lm is a limitation or threshold limit, namely an area that is at risk for development (ha), La is a natural limitation, namely a protected and disaster-prone area and unsuitable soil and hydrological conditions (ha), Lb is the built-in limitation, namely the land use area for non-agricultural cultivation (ha) and fertile agriculture, and infrastructure and utility limitations, namely the area that has been used for infrastructure and utility development in the new city area (ha). Furthermore, the allocation of space utilization and land capability are used as indicators for determining the carrying capacity of the land. To maintain the preservation of the protected function of the area using a coefficient value between 0.3–0.4, which is assumed that 30% is used for the protection function. The formulation used, as follows:

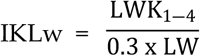

Where IKLw is an index of land capability for new city areas, LWK_1-4_ is the area that has land capability I-IV, LW is the area of the new city area, 0.3 is the coefficient of at least 30% of the protected function of an area (for developing areas), while for undeveloped areas use an index value of 0.4 or more. Furthermore, the carrying capacity is calculated based on the comparison of water availability and demand. The availability and demand for water in the new city area is calculated using the following formulation:

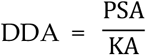

Where DDA is the carrying capacity of water resources, PSA is the potential of water resources, and KA is the population’s clean water needs. Furthermore, the component of the carrying capacity of water resources in terms of water availability uses the following formulation:

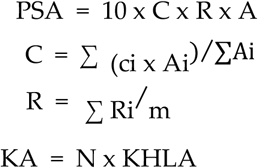

Where PSA is water availability (m3/year), C is the weighted runoff coefficient, Ci is the coefficient of runoff from land use i, Ai is the area of land use i (ha), R is the algebraic average of annual rainfall (mm/year), Ri is the annual rainfall at station i, m is the number of rainfall observation stations, A is the area of the new city area (ha) the value 10 is the conversion factor from mm/ha to m3, KA is the total water demand (m3/year), N is the total population, and KHLA is the water requirement for a decent life. Furthermore, the carrying capacity for a protected function is the ability of a new city area with various land use activities in it. The coefficient number used refers to land use or land cover data. The formulation used in the analysis is as follows:

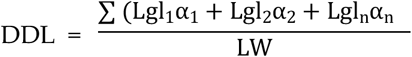

Where DDL is the carrying capacity of the protective function, Lgl_1_ is the area of land use type i, 1 is the coefficient of protection for land use i, and LW is the area of the new city area (ha). The carrying capacity of the protection function (DDL), has a value ranging from 0 (minimum) to 1 (maximum). That is, the closer the value to 1, the better the protection function, and vice versa, if it is close to 0, then the protection function is getting worse. Furthermore, the protected area utilization index is calculated using the following formulation:

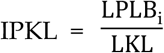

Where IPKL is the index of protected use, LKL is the area of protected areas, and LPLBi is the area of certain cultivated land use (i). Furthermore, the green open space index is calculated using the following formula:

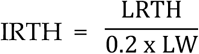

Where IRTH is the index of green open space, LRTH is the area of green open space, and “LW” is the area of the new city area, and 0.2 is the coefficient value of at least 20% of the specified green open space. Furthermore, the determination of the size of the ecological footprint uses the following formulation:

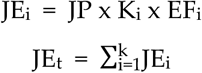

Where JE_i_ is the value of the ecological footprint for land use i (ha), JP is the number of inhabitants (people), K_i_ is the value of land requirement i, to meet the consumption needs of the population per capita (ha/capita), EF_i_ is the equivalence factor and JE_i_ is the total ecological footprint value. Furthermore, the determination of the size of the ecological footprint of bio-capacity using the following formulation:

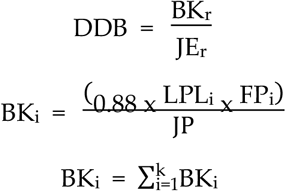

Where DDB is the carrying capacity of bio-ecology, BK_r_ is the total bio-capacity (ha/capita), JE_r_ is the total ecological footprint value. BK_i_ land use bio-capacity i (ha), LPLi land use area i (ha), 0.88 is a constant (12% of which is used to ensure the sustainability of biodiversity, FP_i_ is the factor of production and “JP” is the total population (people). Notes: (1) if DDB value>1, it means BK_r_ > JE_r_ in a condition of surplus, where the ecosystem is able to support the people who live in it = BK_r_ — JE_r_ > 0, (2) if DDB<1, means BK_r_< JE_r_ that there is an overshoot condition, where the ecosystem is not able to support the people who live = BK_r_ — JE_r_< 0. Furthermore, optimal land and population requirements use the following formulation:

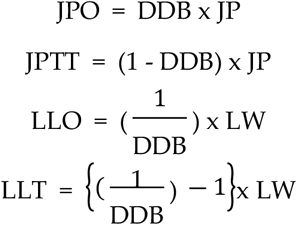

Where “JPO” is the number of residents who are able to be supported by the new urban area ecosystem, “JPTT” is the number of residents who are unable to be supported by the new urban area ecosystem, “LLO” is the optimal land area to support the population in the ecosystem. “LLT “ is the additional land area to support the population in the new urban area ecosystem. Furthermore, DDB is the bioecological carrying capacity, JP is the total population and LW is the area of the new city area.

#### Statistic Analysis

Multiple linear regression method was used to answer research questions, namely the effect of land use change, housing development, infrastructure development, and land reclamation on environmental degradation in the new city area. The measured variables, namely X_1_ (change of land use), X_2_ (residential development), X_3_ (infrastructure development), and X_4_ (land reclamation) to Y (environmental degradation of new city areas). Test the effect between variables using multiple linear regression and correlation test with the following equation:

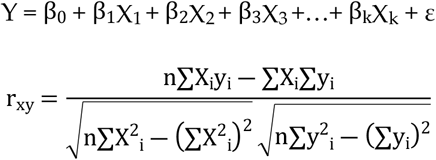

Where Y is the dependent variable, X_1_, X_2_, X_3_, X_4_,,,,, X_k_ is the independent variable, ε is the residue, β_0_, β_1_, β_2_, β_3_, β_4_,,,,,,,,β_k_ is a population parameter whose value is unknown which is estimated from the data. Value β_1_ declare the contribution of the independent variable X_1_ to the dependent variable Y. _rxy_ is the correlation coefficient, n is the number of respondents, ∑X is the score of items, ∑y is the total of the total scores obtained for each respondent, ∑X^2^ is the number of squares of items, ∑y^2^ is the total of the squares of the total scores obtained by each respondent, ∑Xy is the number of multiplication results between the scores of the questionnaire items and the total scores obtained from respondents.

#### SWOT Analysis

SWOT analysis in this study is used to formulate strategies for environmental management and sustainable development of new city areas. The identification and formulation of strategies refers to several considerations, namely strengths, weaknesses, opportunities and threats. The resulting strategy formulation is in the form of the following matrix.

**Table 2.** Swot analysis matrix.

| Internal Factors | External Factors |  |  |
| --- | --- | --- | --- |
|  | Identification of Factor | Opportunity (O) | Threath (T) |
|  |  | Determine the probability factors | Determine the threat factors |
|  | Strenght (S) | S vs. O | S vs. T |
|  | Determine the power factors | Determine the program that emerges by matching the strength (S) with the opportunity (O) | Define emerging programs by matching Strengths with threats |
|  | **Weakness (W)** | **W vs. O** | **W vs. T** |
|  | Determine the weakness factors | Determine the program that emerges by bringing together weaknesses (W) with Opportunities (O) | Determine the program that emerges by meeting weaknesses (W) with threats (T) |

**Source:** Reference [25].

## Results and Discussion

### Determinants of Land Use Change in the New City Area

The dynamics of the new city area development contributes to changes in the characteristics of rural areas and the Mamminasata Metropolitan urban transportation system. Thus, rural areas that are in contact with big cities will experience significant spatial differentiation, socio-economic dynamics and environmental changes [26]. Furthermore, the increase in socio-economic activities coupled with infrastructure development contributes to changes in spatial structure and urban spatial patterns. This means that the centralization of economic activity through the support of urban infrastructure will form a hierarchy of service centers and influence land use patterns [27,28]. The impact of changes in land use in the new city area is presented in Table 3 below.

**Table 3.** The impact of land use change in the new city area of Moncongloe–Pattalassang on the Mamminasata Metropolitan urban system

| Parameter | Impact | Interpretation |
| --- | --- | --- |
| Spatial Structure | Establishment of a hierarchy of service centers and ecosystem complexity | Changes in the urban service structure of the Mamminasata Metropolitan.<br>Polarization of socio-economic activities towards new urban areas and the establishment of growth centers. |
| Spatial Pattern | Agricultural land conversion and decline in agricultural production | Changing the orientation of the activities of rural farming communities to urban industries.<br>Changes in the typology of community housing towards modern architectural housing. |
| Transportation System | Increased traffic volume and traffic jams on main roads | Mobility of freight and passenger transport.<br>Use of private cars, motor vehicles, circular migration and air quality pollution. |
| Ecosystem Condition | Environmental pollution (soil, water and air) | Environmental degradation and reduction of Green Area Coefficient (GAC) and biodiversity.<br>Utilization of water catchment areas and land elevation differences. |
| Population Mubility | Travel generation and traffic pull | Increased population mobility based on the pattern of origin and destination of travel.<br>Increasing the number of movements based on space utilization zones. |
| Economic Activities | Dualistic economics (formal-informal) | Economic inequality and a dualistic economic system develop on the main road corridor.<br>Changes in the traditional community work system towards an urban industrial community. |
| Land Value | Changes in the value of tax objects and land prices | • Changes in the selling value of land based on the function of the developing space.<br>• The economic value of land is determined by the function of commercial space and the market economy. |
| Social Structure | Changes in social interaction and adaptation | • Social relations develop towards economic relations and sharpening of socio-economic stratification.<br>• Changes in the social status of the community based on educational background, expertise, and skills. |

**Source:** Primary data and analysis results

Three things can be explained regarding the results of Table 3, among others: First, the establishment of a hierarchy of urban service centers coupled with changes in spatial patterns contributed positively to the increase in land buying and selling transactions carried out by the community to developers. Furthermore, the spatial dynamics of the new urban area cause changes in the status and value of agricultural land use towards urban land use values based on the function of socio-economic activities in suburban areas. That is, the value of land is determined by economic rent based on land use patterns and developed economic activities [29,30]. This process in addition to changing the typology and morphology of rural areas, also contributes to increasing the flow of development investment towards the formation of spatial zoning and community segmentation. That is, the spatial zoning that is built has an impact on the grouping of people based on economic capacity and income level. Urban planning and economic forces are the main drivers of expansion of housing space, income inequality and various socioeconomic problems [31,32]. Thus, the allocation of space and land in suburban areas is created due to the support of government policies and the effects of the development of new city areas. The spatial distribution of suburban areas is related to land characteristics and location characteristics, in the sense that plans are formulated and implemented to accommodate urban development needs [33,34].

Second, the development of new urban areas tends to increase from time to time, which contributes to the increase in population mobility and affects the urban transportation system based on the pattern of origin and destination of travel. This condition is characterized by an increase in traffic volume, circular migration, and the distribution of the flow of goods and services in the direction of decreasing environmental quality. Furthermore, the increase in socio-economic activities coupled with population mobility and the development of transportation infrastructure have an impact on reducing the coefficient of green areas and biodiversity and environmental pollution. Thus, it is necessary to regulate density, diversity, mixed land use, sustainable transportation, and green space through support for strengthening the capacity of government and private institutions as development actors [35]. Third, the dualistic economy that develops in addition to influencing land values, also contributes to changes in the work system of rural communities towards urban industries. These dynamics cause changes in the social structure of society towards differences in status and economic strata based on abilities and expertise as well as changes in people’s lifestyles. Thus, the land value of the new city area is no longer measured based on the area and productivity of agricultural land, but is measured based on the function of developing urban space. Thus, built-up areas in suburban areas have led to a significant decrease in agricultural area, gardens, and vacant land as well as changes in land values [36]. The development activities of the new city area are presented in Figure 2 below.

**Fig. 2.**
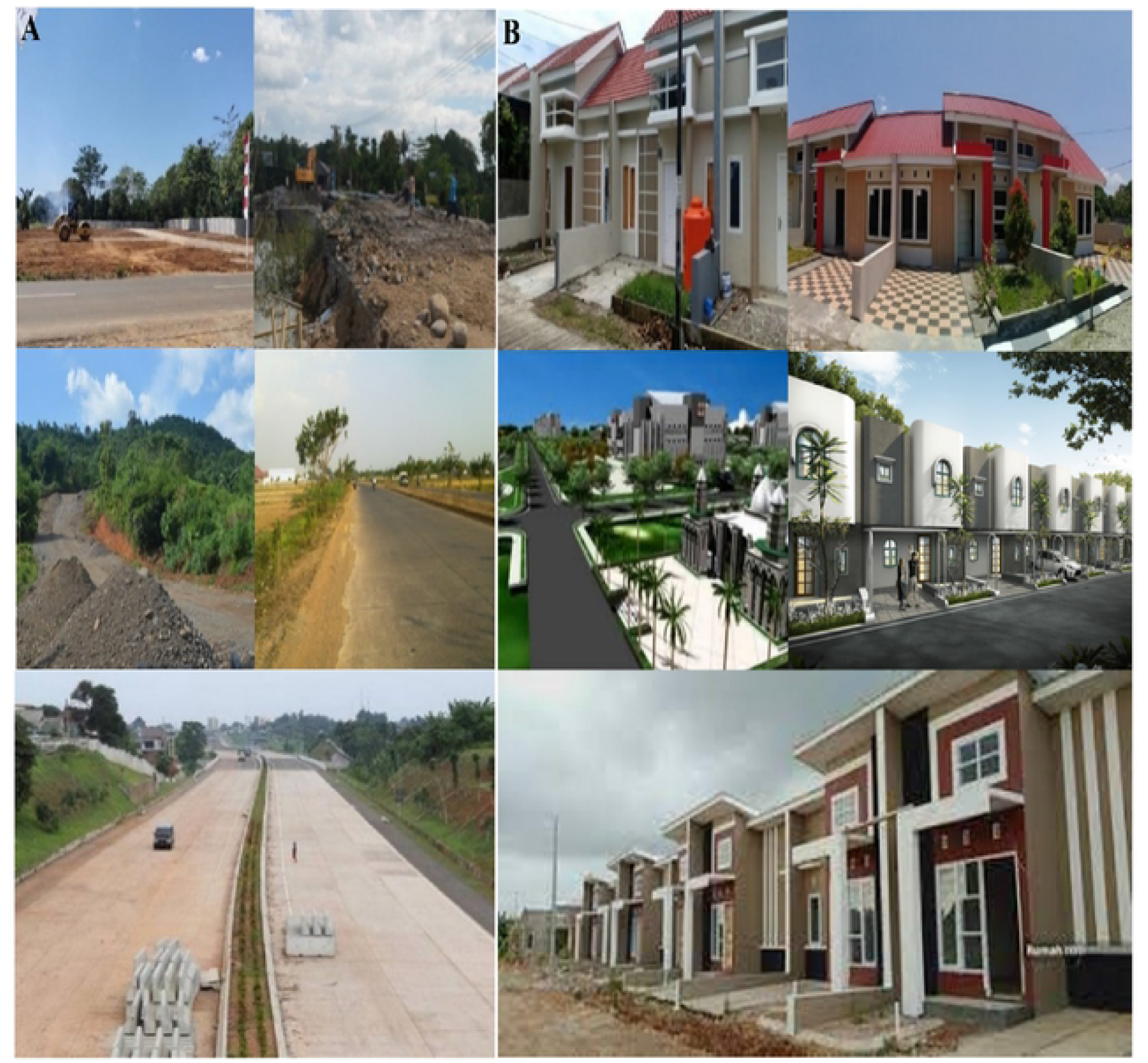
(A–B). Development of the new city area of the Moncongloe-Pattalassang. (A) Land reclamation and road infrastructure development; (B) Housing development and socio-economic activities.

Figure 2A shows the new city area of Moncongloe–Pattalassang built through agricultural land conversion, land clearing, and land reclamation. This process is carried out to meet the needs of housing development, road infrastructure, and the provision of socio-economic facilities. Conversion of agricultural land that tends to increase and the use of water catchment areas for the development of new urban areas has an impact on changes in ecosystem characteristics and the complexity of the urban transportation system. Changes in land use significantly affect the value of ecosystem services (ESV), ecosystem functions and urban resilience reflecting a multi-level and multi-dimensional dynamic accumulation process [37,38]. Thus, government policy support is needed to anticipate environmental damage and the risk of flood threats through disaster mitigation of the impacts [39,40].

Figure 2B, shows that housing development that is being developed towards large-scale settlements has a positive contribution to the complexity of the ecosystem due to land cover changes. This condition has become the driving force for the entry of migrants to occupy elite settlement locations towards the separation of migrant settlements and local communities. This means that the construction of settlements in the new city area has the effect of discrimination, eviction, and displacement on the existence of the local community [41]. There are four categories of change processes that arise due to the intensity of settlement development, namely (i) changes in land use forms, (ii) changes in building characteristics, (iii) residential typology, and (iv) changes in social interaction and social adaptation. These four things trigger the socio-economic dynamics of society towards economic disparity in wealth, segregation, and the emergence of an exclusive society. That is, the process shows a widening income gap and suburban areas are developing towards development oriented towards meeting the needs of certain groups [42]. Changes in land use in the new city area are presented in Figure 3 below.

**Fig. 3.**
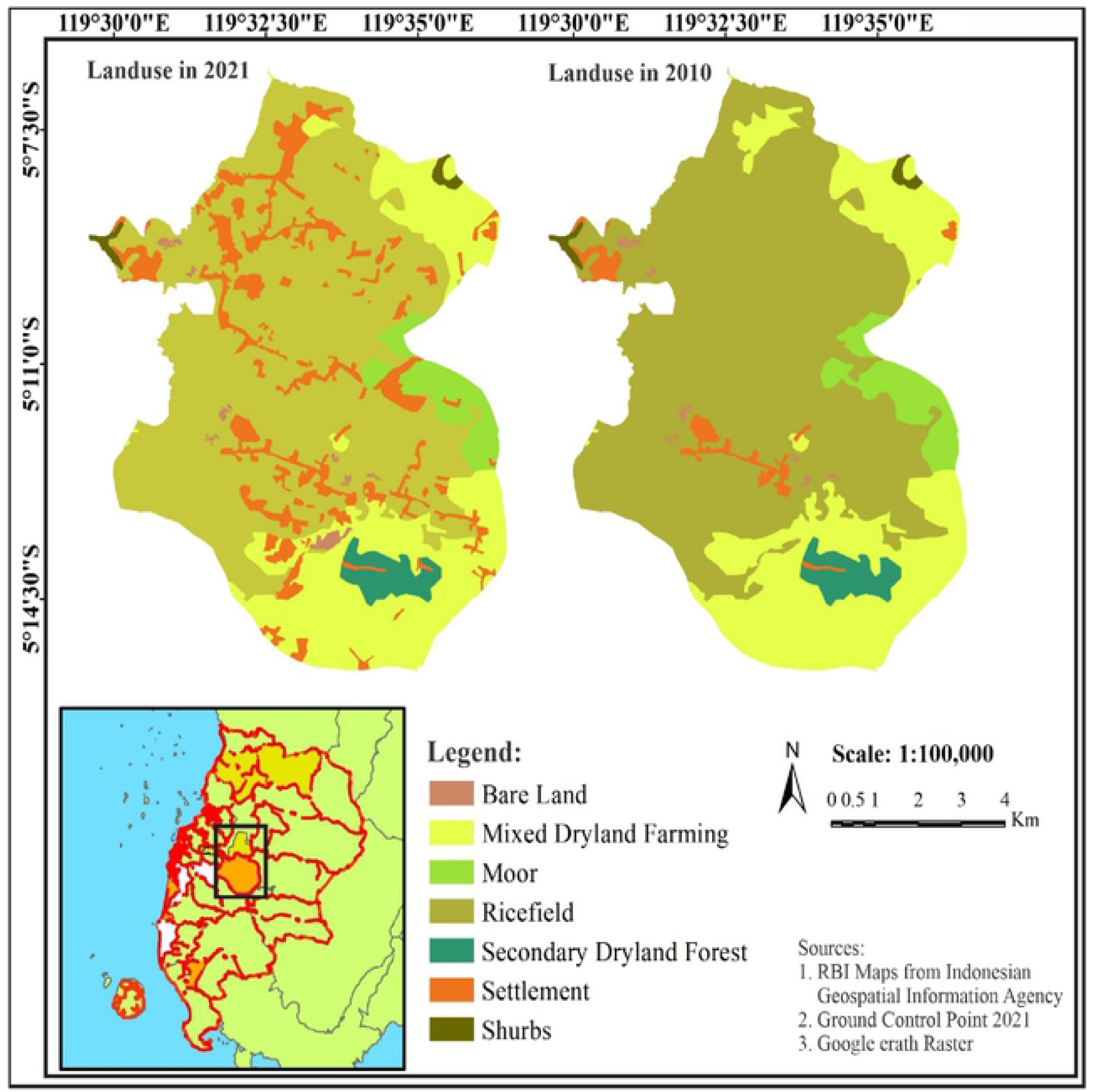
Changes in land use in the new city area of the Moncongloe–Pattalassang

The possible interpretations of Figure 3 include: First, the intensity of land use change is closely related to housing and settlement development. Second, the use of space during the 2010 period was dominant in utilizing productive agricultural land towards horizontal spatial physical development and its influence on changes in the spatial attribution of rural areas as well as changes in the orientation of community activities from traditional agrarian to urban industrial and changes in the structure and pattern of urban space in the Mamminasata Metropolitan area. This means that spatial connectivity has an impact on increasing industrial-oriented population flows and population mobility causing changes to the agricultural system in rural areas [43]. Third, in the period from 2015 to 2021, the expansion of the new city area was then developed to support the construction of the Moncongloe by-pass road infrastructure towards Hasanuddin International Airport. The transportation infrastructure in addition to providing ease of movement and accessibility also has an impact on population mobility, circular migration, and increased traffic volume. This means that transportation activities contribute to water, air, and soil pollution [44]. Thus, transportation system connectivity, in addition to contributing to the integration of urban systems, also contributes to environmental degradation. Road–based transportation has the potential as a source of environmental pollution, economic losses, safety and health risks due to traffic jams [45].

### Decline in the Environmental Quality of the New City Area

The decline in environmental quality is the effect of changes in land use, increased socio-economic activities and infrastructure development. Conversion of productive agricultural land and ecosystems into built–up areas has an impact on ecological conditions, damage to natural resources and natural vegetation in suburban areas [46–48]. The facts found in the field indicate that the sources of pollution in the new city area include: (1) Waste originating from households, shops, education, hospitals, and urban informal economy activities; (2) Waste originating from households, socio-economic activities, and roads; (3) Residential building construction and infrastructure development; (4) Air pollution produced by motor vehicle exhaust gases; and (5) land clearing and land reclamation. Thus, development activities that tend to increase every year are positively associated with ecosystem stability. This means that the imbalance in development will have an impact on changes in the ecosystem, economy, and social life of the community [49,50]. Thus, it is necessary to monitor development actors, protect ecology, and collaborate in environmental management [51]. Sources of environmental pollution are presented in Figure 4 below.

**Fig. 4.**
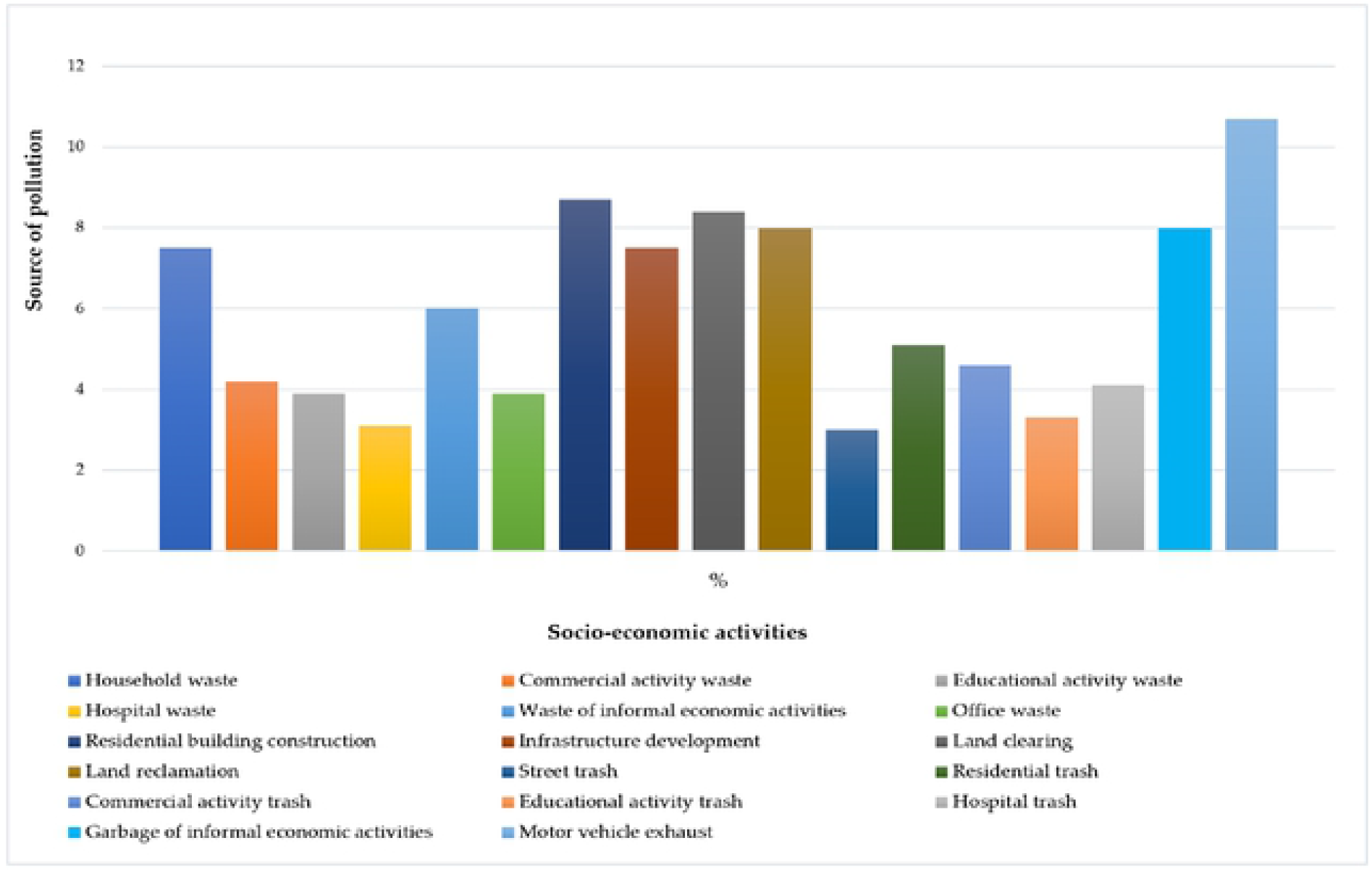
Sources of environmental pollution in the new city area of Moncongloe-Pattalassang. Source: Primary data

The proposed interpretations are related to Figure 4, among others: (1) Domestic waste generated by socioeconomic activities contributes 28.6%; (2) Building construction, infrastructure development, land clearing, and reclamation contributed 32.6%; (3) Domestic waste originating from socio-economic activities contributed 28.1%; and (4) Motor vehicle exhaust gas contributed 10.7%. These results confirm that the development of new urban areas causes changes in land cover, changes in ecosystem characteristics, economics, social and spatial use conflicts in suburban areas. This means that the planning mechanism applied for the development of suburban areas contributes to changes in land use, changes in rural characteristics, social stratification, polarization, and environmental imbalances [52,53]. The scale of development that tends to increase is positively associated with the use of water catchment areas, reduction of forest cover and disturbance to biodiversity. This means that an increase in development activities and human activities is the cause of the potential risk of natural disasters due to damage to environmental ecosystems [54]. The facts found in the field illustrate that forest destruction and the use of water catchment areas contribute to changes in the flood cycle in the suburban area of the Mamminasata Metropolitan. Flood is a natural disaster that causes disease, damage and loss of life, property, and infrastructure as well as disruption of public services [55]. The carrying capacity of the residential environment based on spatial zoning is presented in Figure 4 below.

Figure 5 shows the development of new urban settlements based on spatial zoning. Interpretations that can be put forward to these results include: (1) The development of dominant settlements utilizing productive agricultural land has a direct impact on ecosystem conditions; (2) Coverage of buildings in zones A, B, and E is greater than 70%. This means that the activities developed have exceeded the optimum limit for land use; (3) Thus, the potential area of land that is possible to be utilized to support the development of settlements is 70% of the total area of the new city area. These results confirm that settlement development in addition to contributing to space pressure also affects the provision of green open space by 30%. The construction of settlements on productive agricultural land has a direct impact on the quality of life of the community, water quality pollution and ecological stability [56]. Thus, it is necessary to evaluate the mechanisms and procedures for implementing new city development to ensure ecological sustainability and to anticipate the impacts that will arise and the risk of natural disaster threats. Thus, sustainable development is a very important procedure to be implemented to ensure social, economic and environmental sustainability [57]. The land capability index and the need for clean water services are presented in Figure 6 below.

**Fig.5.**
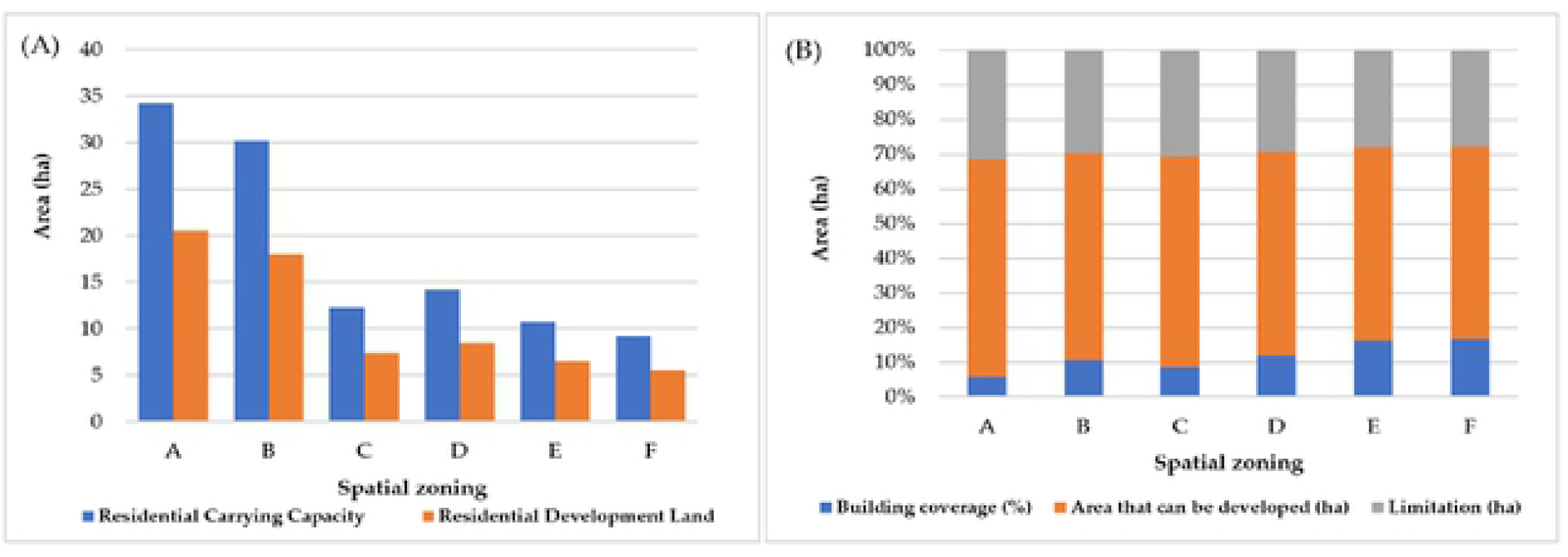
(A–B). Construction of settlements in the new city area of Moncongloe – Pattalassang. (A) Carrying capacity and land for development; (B) Coverage of buildings and development thresholds. Source: Analysis results.

**Fig.6.**
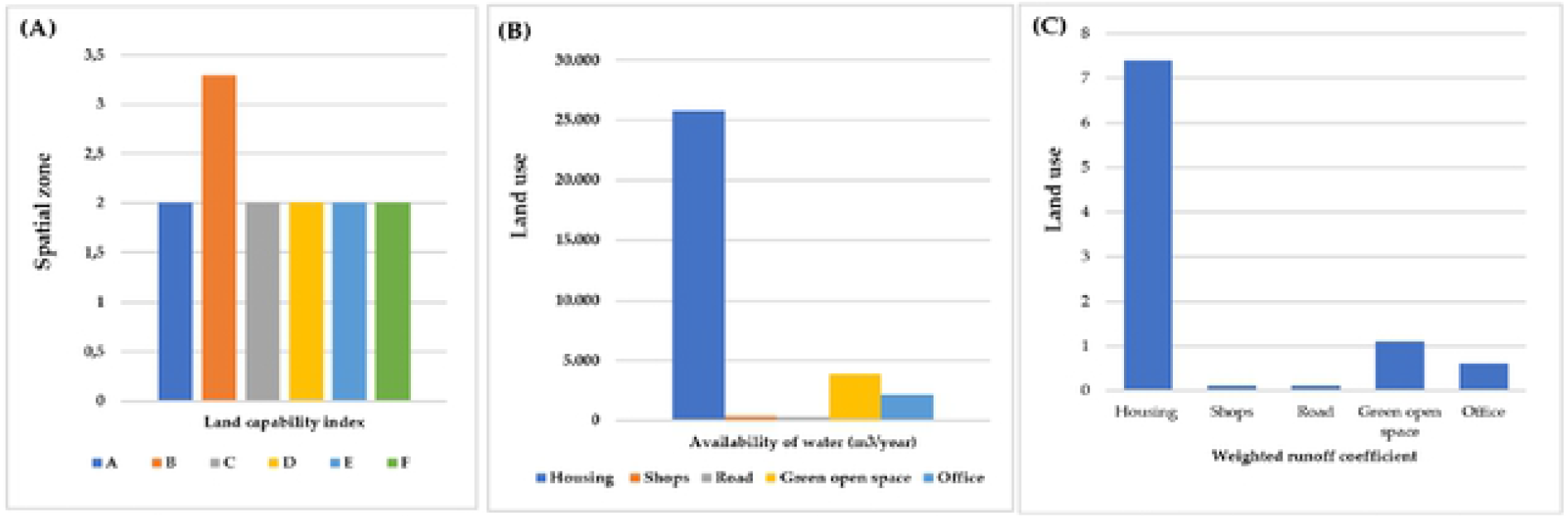
(A–C). Carrying capacity of land and carrying capacity of water resources. (A) Land cover index; (B) Availability of water; (C) Runoff coefficient. Source: Analysis results

Figure 6A can be explained, namely: (1) The highest land capability index is located in zone B with a value of 3.3; and (2) Zones A, C, D, E and F with a land capability index value of 2.0. This figure confirms that the land in the new city area has the potential to support various cultivation activities including for housing and settlement development while ensuring environmental balance. Therefore, it is necessary to evaluate the development of settlements based on controlling the use of space and the carrying capacity of the environment [58]. Figure 6B can be explained, among others: (1) The need for clean water services for housing is 79.42%, 1.08% for shopping needs, and 0.97% for road infrastructure needs; (2) The need for clean water services for green open spaces is 11.91% and 6.62% for office needs. These results confirm that the development of new urban areas will require the support of potential sources of raw water to fulfill the distribution of clean water services for a population of 5,500 families or 27,500 people. Facts found in the field show that limited clean water services are one of the factors that cause people to use deep groundwater wells with a depth of between 10-20 meters. The use of groundwater has an impact on three things, namely (i) land subsidence due to excessive and continuous use of groundwater, (ii) intensive use of water causes the water quality to be unfit for consumption, and (iii) continuous use of groundwater risk to the slope of buildings and roads as well as public safety. These three things have the potential to threaten the sustainability and balance of the environment. Overuse of groundwater causes ground surface movement, significant and repeated damage to infrastructure and aquifer water storage capacity [59].

Figure 6C can be explained, among others: (1) The highest runoff is generated by housing with a weighted coefficient value of 7.4; (2) The weighted runoff contributed by shops and roads is 0.1, offices are 0.6 and 1.1 is contributed by open space with a drainage area of 4,300 ha. These results confirm that 70% of the rain that falls is channeled into surface runoff, while 30% is lost due to infiltration and evapotranspiration processes, thus potentially causing inundation and urban flooding. Furthermore, urban flooding is influenced by several factors, namely the drainage system, runoff coefficient and land elevation [60]. The facts found indicate that changes in land use have an influence on changes in the hydrological cycle, damage to natural vegetation and changes in forest cover. Thus, making infiltration wells is very important to be held and placed in strategic locations. Furthermore, each residential building requires the support of making biophores with a diameter of 10–30 cm with a depth of 80–100 meters which functions to absorb water and prevent flooding. Thus, Blue and Green Infrastructure (BGI) is an important, nature–friendly measure to manage urban flood risk [61]. The protected area utilization index is presented in Figure 7 below.

**Fig.7.**
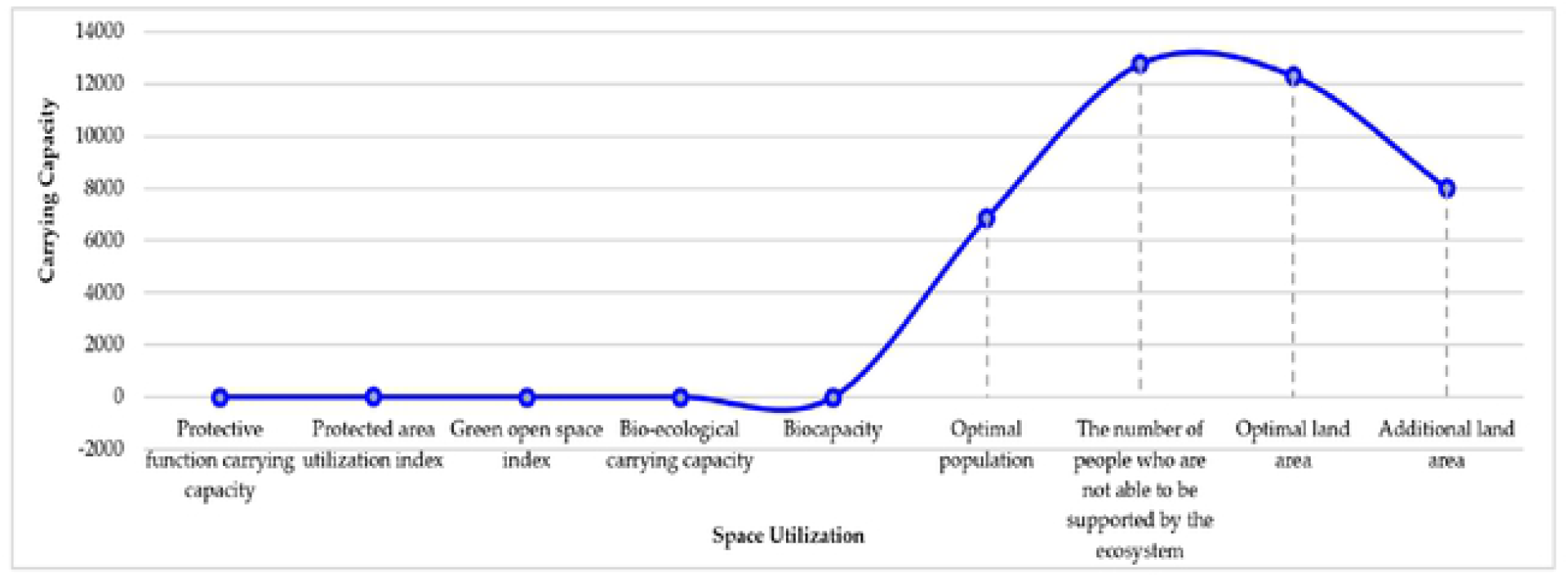
Index of the use of protected areas and the carrying capacity of the new city area of Moncongloe-Pattalassang. Source: Analysis results

The possible interpretations of Figure 7 include: (1) The carrying capacity of the protected function is obtained with a value of 0.35 and an index of utilization of protected areas of 1.1. This figure confirms that the protected function of the new city area is categorized as poor; (2) The green open space index value shows a number of 0.3. This means that the availability of green open space is less than optimal in its function in protecting ecosystem sustainability; and (3) the carrying capacity of bio-ecology obtained a value of 0.8. This means that the condition of the ecosystem does not support the existence of the people who live. Thus, it can be concluded that the carrying capacity and function of the forest is less than optimal due to the complexity of space use and the availability of land for green open space which has shifted its function to the needs of developing socio-economic activities and housing development. Therefore, the expansion of urban land use will have an impact on changes in landscape characteristics, environmental carrying capacity and ecological vulnerability [62]. Furthermore, the results of the survey for the development of new city areas are presented in Figure 8 below.

**Fig.8.**
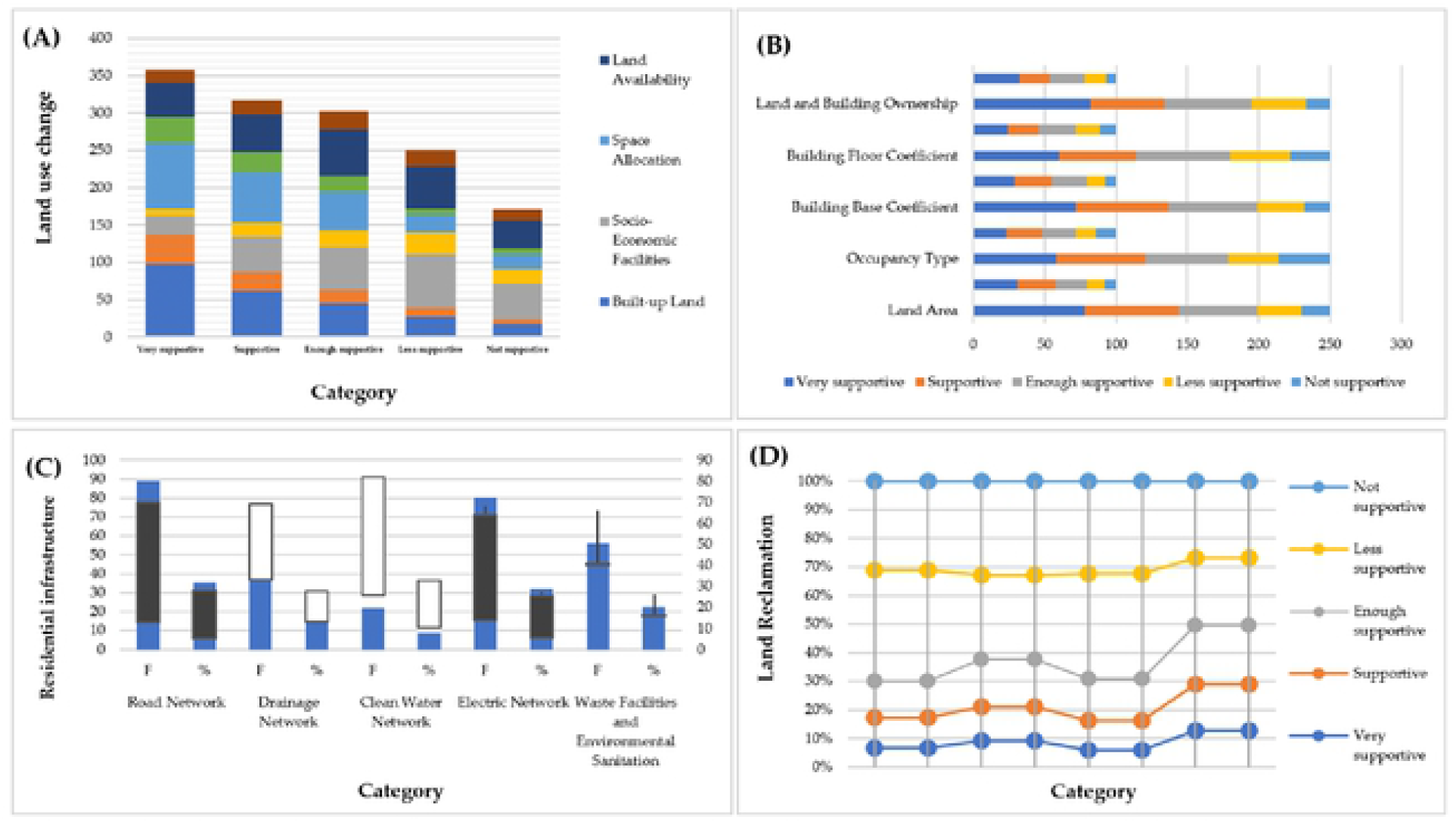
(A–D). Construction of the new city area Moncongloe–Pattalassang. (A) Changes in land use; (B) Fulfillment of housing needs; (C) Fulfillment of infrastructure needs; (D) Land reclamation. Source: Primary data

Figure 8A can be explained, among others: First, the built-up land depicts 64% in the supportive category, 18% enough supportive category and 18% not supportive category. These results indicate that the dominant area of the new city area used is the result of land reclamation of forest areas and water catchment areas carried out by developers through the land clearing process to accommodate housing development, infrastructure, and other socio-economic activities. This process causes land cover changes, ecosystem damage, and a decrease in the carrying capacity of the land. Second, the fulfillment of socio-economic facilities to serve the needs of the settlers, obtained an illustration of 29.2% for the supportive category, 22.8% enough supportive category, and 48% not supportive category. These results confirm that the availability of socio-economic facilities in terms of service coverage has not been optimal in supporting residents who live in the new city area. Third, the need for space to accommodate development needs is shown at 62% for the supportive category, 20.8% enough supportive category, and 17.2% not supportive category. These results confirm that the opportunities for developing new urban areas are influenced by limiting factors, namely forest areas, watersheds, and productive agricultural areas. That is, the process of urban expansion through the construction of new cities has an impact on the depletion of agricultural land and forest destruction leading to environmental damage and land carrying capacity [63,64]. Fourth, the availability of land obtained a value of 38% for the supportive category, 24.8% enough supportive category, and 37.2% not supportive category. These results confirm that the availability of land and the environmental carrying capacity of the new city area is quite low in the sense that it requires monitoring and control of land use in the future. Thus, it is very important to implement a compact city design in the development of new city areas to ensure environmental, economic and social sustainability [34,65].

Figure 8B which can be explained, among others: (1) The area of land development for housing obtained an overview of 57.6% in the category of supportive, 22% enough supportive, and 20.4% not supportive. This figure shows that it is necessary to strictly limit housing development and avoid land use that is not in accordance with the established land use. That is, it is necessary to integrate zoning boundaries by taking into account the spatial heterogeneity associated with population and building density [66]. (2) The type of housing developed is 48.4% in the supportive category, 23.2% enough supportive category, and 28.4% not supportive category. These results illustrate that some of the housing built does not meet the expectations of the community in terms of building construction and the materials used. (3) The basic building coefficient shows 54.8% in the supportive category, 24.8% enough supportive category, and 20.4% not supportive category. (4) The building floor coefficient is 45.6% for the supportive category, 26.4% enough supportive category, and 28% not supportive category. These results confirm that efforts are needed to regulate the area of residential buildings that function to cover the ground surface, to ensure groundwater infiltration or the availability of ground water and to regulate the ground surface that is not covered by buildings to receive direct sunlight so that the soil can dry out and the air created in around the building does not become damp. This means that the handling is needed to anticipate water runoff during high-intensity rains as well as to reduce the burden on the drainage system [67]. Land ownership obtained a value of 53.6% for the supportive category, 24.4% enough supportive, and 22% not supportive category. The lift confirmed that consistent efforts and land tenure arrangements are needed to ensure the sustainability of the urban ecosystem. Furthermore, the development of ecological networks is an important element to reduce the degradation of urban habitats and protect the natural environment [68].

Figure 8C can be explained: (1) The road network system shows 63.6% in the supportive category, 22.4% enough supportive, and 14% not supportive category. This figure illustrates that the quality of the road network still requires quality improvement. (1) The environmental drainage network is 31.6% supportive, 17.6% enough supportive, and 58.8% not supportive. The facts found in the field illustrate that the function of the drainage channel in the new city area which is less than optimal in draining rainwater is positively associated with urban flooding that occurs every year. Urban flooding is influenced by three factors, namely the capacity of the drainage network system, environmental changes and reduced watershed permeability due to urban growth [69]. (3) The fulfillment of clean water services is 19.2% supportive, 16.4% enough supportive, and 64.4% not supportive. The appointment confirmed that the distribution of clean water in the new city area had not been fulfilled and did not meet the minimum service standards. (4) Fulfillment of electrical energy with a value of 57.6% for the supportive category, 27.2% enough supportive, and 15.2% not supportive. This figure shows that electrical energy services have not been fulfilled optimally. (5) The availability of solid waste facilities and environmental sanitation shows a score of 38.4% for the supportive category, 18.8% enough supportive, and 42.8% not supportive. This figure confirms that the domestic waste produced is not managed optimally and that environmental sanitation has not been fulfilled is one of the factors that causes environmental pollution in the new city area of Moncongloe–Pattalassang. This means that waste management and poor sanitation are factors that cause various environmental and social impacts and affect public health [70].

Figure 8D that can be explained, among others: (1) Land elevation obtained a value of 17.2% for the supportive category, 12.8% enough supportive, and 70% not supportive category. This figure shows that the variation of land elevation for each zone and different housing blocks is positively associated with waterlogging and a fairly high runoff coefficient and is a factor that causes flooding in the residential environment. (2) The development threshold is 21.2% for the supportive category, 16.4% enough supportive, and 62.4% not supportive category. These results confirm that efforts to limit housing development are needed to anticipate ecosystem damage and in particular to secure river borders and water catchment areas. (3) Mechanisms and procedures for land reclamation in the new urban area, the figure is 16.4% in the supportive category, 36.8% enough supportive, and 69.2% not supportive category. These results confirm that the land reclamation carried out has not referred to the standard provisions of land use and ecosystem revegetation that have been set by the government. Thus, efforts to optimize land use and ecosystem revegetation are needed for the purpose of creating a balance between socio-economic systems and natural ecosystems [71]. (4) The carrying capacity of the land shows a value of 28.8% in the supportive category, 20.8% enough supportive, and 50.4% not supportive category. These results illustrate that the development of new urban areas has exceeded the carrying capacity of the environment in relation to the availability of land and the number of residents that can be accommodated. Thus, efforts are needed to integrate the pattern of use with the city activity system towards development [72,73]. Furthermore, the effect of land use change, housing development, infrastructure development, and land reclamation on environmental degradation in the new city area is presented in Table 4 below.

**Table 4.**
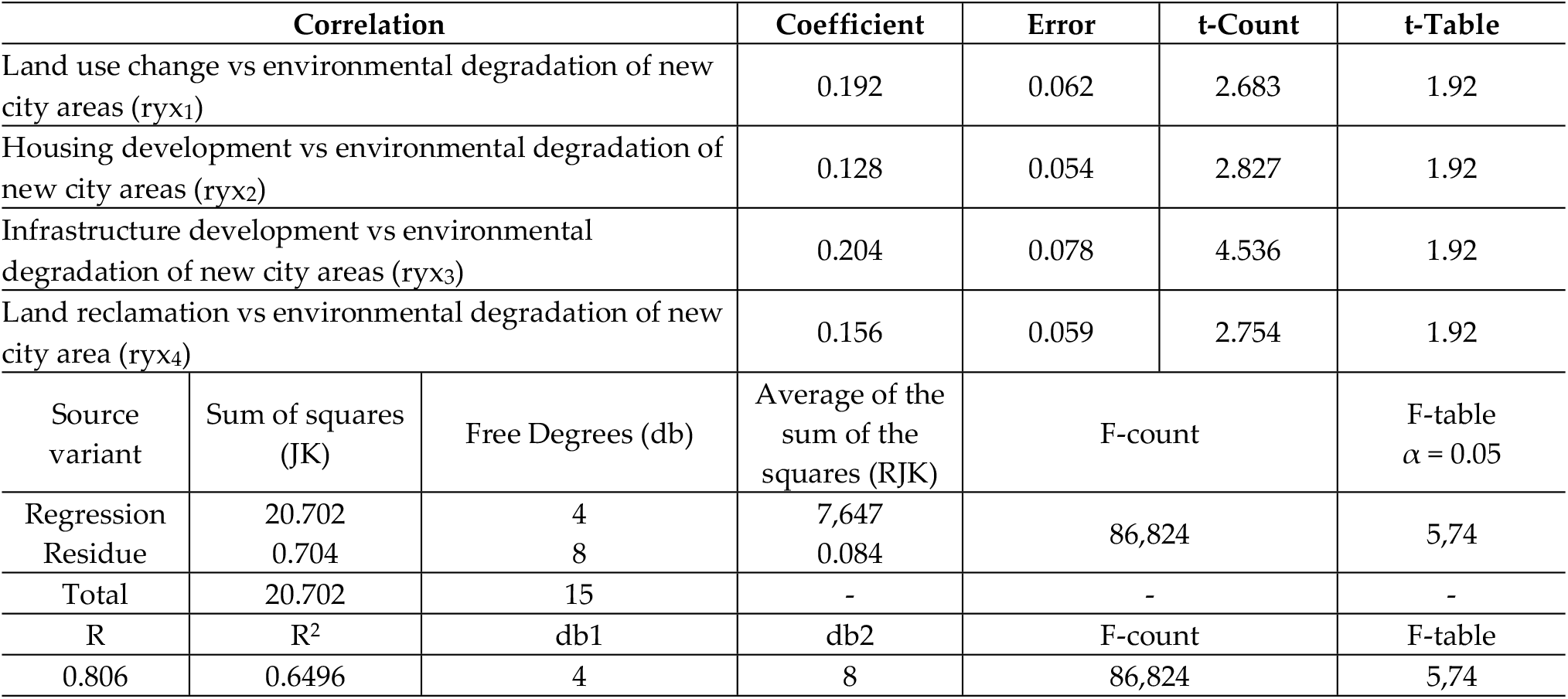
Summary of the results of the regression coefficient significance test.

| Correlation |  |  | Coefficient | Error | t-Count | t-Table |
| --- | --- | --- | --- | --- | --- | --- |
| Land use change vs environmental degradation of new city areas ($ryx_1$) | | | 0.192 | 0.062 | 2.683 | 1.92 |
| Housing development vs environmental degradation of new city areas ($ryx_2$) | | | 0.128 | 0.054 | 2.827 | 1.92 |
| Infrastructure development vs environmental degradation of new city areas ($ryx_3$) | | | 0.204 | 0.078 | 4.536 | 1.92 |
| Land reclamation vs environmental degradation of new city area ($ryx_4$) | | | 0.156 | 0.059 | 2.754 | 1.92 |
| Source variant | Sum of squares (JK) | Free Degrees (db) | Average of the sum of the squares (RJK) | F-count | | F-table $\alpha = 0.05$ |
| Regression | 20.702 | 4 | 7,647 | 86,824 |  | 5,74 |
| Residue | 0.704 | 8 | 0.084 |  |  |  |
| Total | 20.702 | 15 | - | - |  | - |
| R | R<sup>2</sup> | db1 | db2 | F-count |  | F-table |
| 0.806 | 0.6496 | 4 | 8 | 86,824 |  | 5,74 |

The possible interpretations of the results of Table 4, namely: (1) Changes in land use have a positive effect on environmental degradation in the new city area; (2) Housing development has a positive effect on environmental degradation in the new city area; (3) Infrastructure development has a positive effect on environmental degradation of the new city area; and (4) land reclamation has a positive effect on environmental degradation in the new city area. These results confirm that changes in land use, housing development, infrastructure development, and land reclamation explain 64.96% of the environmental degradation of the new city area of Mongcongloe–Pattalassang. This means that the intensity of land use changes due to development activities coupled with climate change causes land degradation [74,75]. Thus, the demands of urban development today are very important and strategic to be oriented towards sustainable development [76,77].

### Strategy for Environmental Management and Sustainability of New City Area Development

The spatial dynamics and integration of the Mamminasata urban system is part of the impact of the development of the new city area which requires handling and control actions towards the implementation of sustainable development. An integrated and holistic approach is needed to build urban resilience through policy formulation towards the development of green infrastructure to overcome ecosystem disturbances and natural disasters [78,79]. Furthermore, the strategic steps needed to support the formulation of government policies are presented in the following Table 5 matrix.

**Table 5.** Strategy for improving environmental quality and sustainable development of the new city area of Moncongloe–Pattalassang.

| External | Internal | Strength | Weakness |
| --- | --- | --- | --- |
|  |  | The area for the development of the new city area is sufficient.<br>Government policy support. | Conversion of productive agricultural land. |

|  | • The growth pole of the Mamminasata Metropolitan area.<br>• Regional connectivity and strategic location.<br>• Urban transport network system connectivity. | • Environmental carrying capacity is quite low.<br>• Urban flooding.<br>• Environmental damage and conversion of forest area functions.<br>• Land reclamation and reduction of water catchment areas. |
| --- | --- | --- |
| Opportunity | Strategy (SO) | Strategy (WO) |
| • Sufficient investment support.<br>• Demand for housing needs is quite high.<br>• Implementasi konsep compact city.<br>• Transportation movement system is quite high.<br>• Large-scale settlement development. | • Optimizing the utilization of development investment towards the effectiveness and efficiency of land use and saving the environment.<br>• Implementation of government policies towards fulfilling livable housing standards and based on natural disaster mitigation.<br>• Development of new urban areas as new growth poles through the implementation of the compact city concept based on mixed land use patterns.<br>• Management of the transportation system through the support of green infrastructure and green building.<br>• Development of large-scale settlements coupled with improving the quality of infrastructure and optimal road performance. | • Optimizing the utilization of development investment and minimizing conversion of productive agricultural land.<br>• Limiting the development of new settlements in water catchment areas and improving environmental quality.<br>• Implementation of the concept of a compact city based on a balanced housing pattern.<br>• Limiting the development of transportation infrastructure in forest area locations.<br>• Development of large-scale settlements based on natural disaster mitigation and environmental sustainability. |
| Threats | Strategy (WO) | Strategy (WT) |
| • Climate change.<br>• Potential for natural disasters.<br>• Environmental pollution<br>• Space use conflicts<br>• Intensity of land use change and changes in environmental situation | • Adaptation to climate change towards environmental carrying capacity resilience.<br>• Mitigation of natural disasters by providing evacuation facilities in strategic locations and easily accessible by residents.<br>• Land use administration arrangement based on land management application.<br>• Strict land use regulation based on zoning arrangements for new urban areas.<br>• Development of a mass transit-based transportation network system. | • Adaptation to climate change based on food security and sustainable energy.<br>• Improvement of environmental quality based on natural disaster mitigation.<br>• Control of environmental pollution and restoration of environmental quality.<br>• Conservation of forest areas and management of spatial use conflicts.<br>• Implementation of standard provisions for land use and ecosystem revegetation in new city areas. |

The results of the strategy formulation (Table 5) which can be explained include: (1) The implementation of strategic policies for the development of new urban areas requires the involvement of various parties as development implementers; (2) The development of new urban areas requires inter-sectoral and cross-sectoral synergy; (3) Development of settlements and support for infrastructure development as well as other activities oriented towards saving the environment; and (4) the balance of development is carried out through monitoring and supervision based on the participation of stakeholders. The results of the strategic formulation will require cooperation between the government, the private sector, and the community in realizing the sustainable development of the new city area of Moncongloe–Pattalassang. Strategic policy steps and the SDGs governance system will require the cooperation of various parties, private and community support towards the sustainable development of new city areas [80–82]. The sustainability of the development of the new city area as an integral part of the Mamminasata Metropolitan urban system is presented in Figure 9 below.

**Fig.9.**
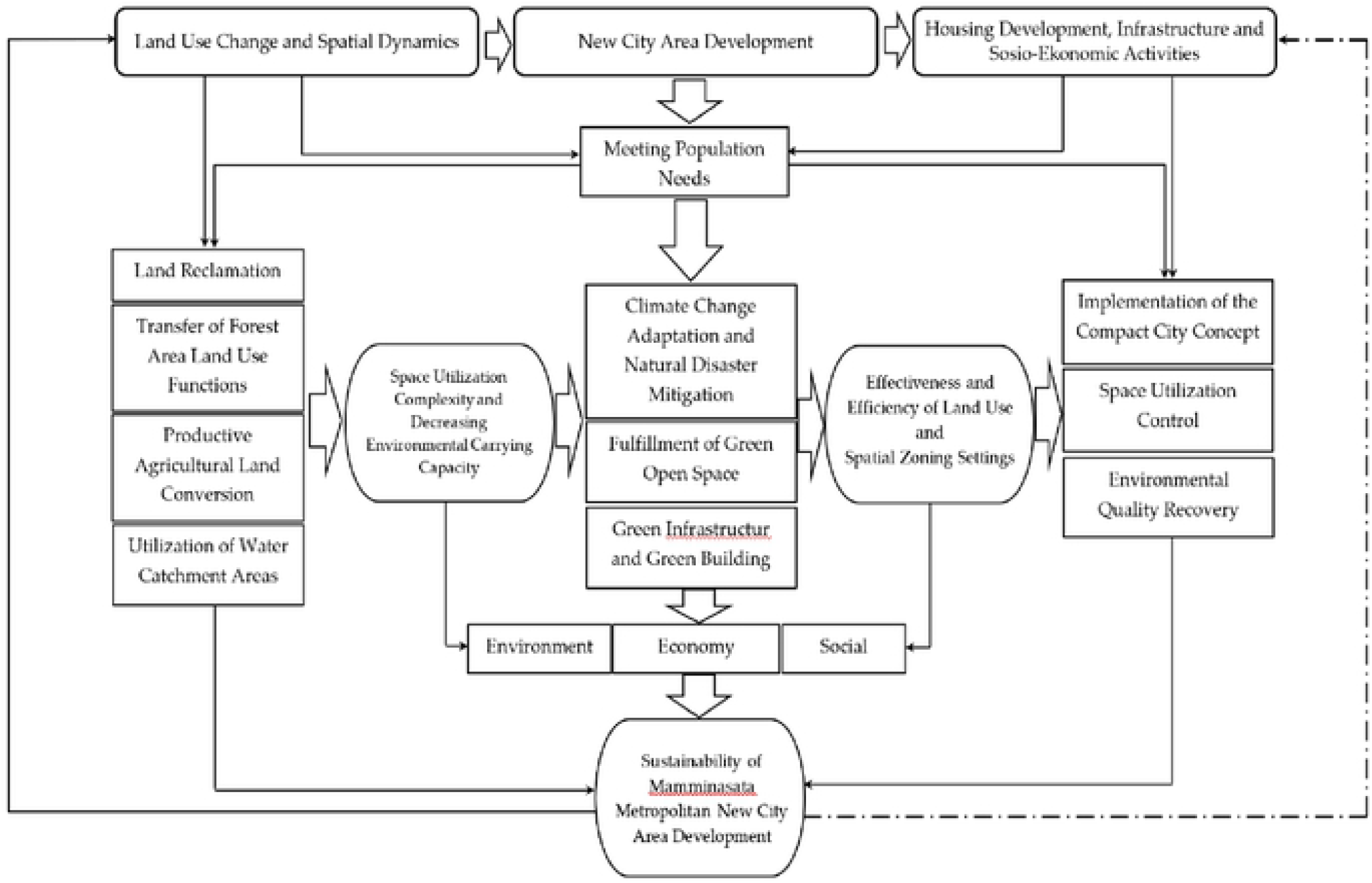
Sustainability of development of the new city area of Mamminasata Metropolitan. Source: Author’s elaboration

Interpretations of Figure 9 that can be proposed include: First, changes in land use and spatial dynamics of new urban areas in relation to housing development, infrastructure and social activities through land reclamation have an impact on conversion of productive agricultural land and land use conversion of forest areas. This process contributes to the complexity of space utilization and a decrease in the carrying capacity of the environment. Thus, the strategic actions needed to deal with this, namely (i) tightening the use of water catchment areas, (ii) regulating building boundaries, road borders, road–owned areas and road monitoring areas strictly, (iii) limiting housing development and horizontal settlements towards a vertical development pattern, (iv) integrating balanced housing and settlement development through adequate infrastructure support, (v) limiting development to productive land and forest areas through zoning arrangements and implementing incentives. These five things will create a balance of development towards the sustainability of the ecosystem of the new city area. Thus, efforts are needed to redesign and restructure the urban environment by adopting innovative solutions towards sustainable development [83].

Second, climate change adaptation and natural disaster mitigation are implemented through two main principles, namely (i) providing green open space, and (ii) developing green infrastructure and green building. These two things require the implementation of strategic policies, including: (1) Fulfillment of green open space by 30% of the total area of the new city area; (2) Acquisition of green open space by applying the Green Basic Coefficient (GBC) on private land owned by the public and the private sector; (3) Refunctioning of existing green open spaces through revitalization of forest areas and planting grass in residential neighborhood parks; (4) Utilization of the built space through planting plants on the roof or walls of the building; (5) Developing city green space corridors through the concept of urban park connectors that are connected to other green open spaces; and (6) Establish a consistent location that serves as a water retention pond. Third, the implementation of the compact city concept is oriented towards mixed land use by considering three important aspects, namely (i) saving the environment, (ii) implementing the green economy concept, and (iii) involving community participation in environmental monitoring. These three things will be achieved through strategic synergy by developing environmentally friendly cities towards sustainable development [84].

In the future, the sustainability of new urban areas will be developed towards the implementation of the concept of environmental sustainability, namely: First, the effectiveness and efficiency of land use through consistent development based on the principles of environmental sustainability and natural disaster mitigation. This principle is implemented through several strategic steps, including: (1) Replacing plant varieties of vegetation that are resistant to extreme temperatures and capable of absorbing carbon dioxide and are placed on main roads in the new city area; (2) Ensuring the sustainability of forest ecosystems and essential ecosystems due to the impact of climate change and maintaining biodiversity and ecosystem services to remain sustainable; (3) Optimizing the utilization of organic waste and utilization of biofuel energy sources (new and renewable energy); (4) Building public awareness to maintain a clean environment and water storage to prevent the spread of disease through a strict monitoring system; (5) Managing the environment of the new city area in a sustainable manner; and (6) Strengthening government and community institutional capacity. Thus, a strategic approach is needed to create a city development management system towards the development of sustainable green infrastructure [85,86].

The direction of economic sustainability is oriented to several principles, including: (1) Creating community economic competitiveness; (2) Preparation of decent employment opportunities and encouraging the improvement of strategic economic performance by supporting the improvement of basic infrastructure performance; (3) Minimizing economic and social disparities through creating equal opportunities for the community; (4) Implementation of integrated development through mixed land use as much as possible, ensuring the sustainability of green open space and transportation as a unified system. Furthermore, the direction of social sustainability is oriented in several principles, namely: (1) Creation of intergenerational equality and justice; (2) Provision of optimal social services through health services, education, gender equality, and public accountability; (3) Guarantee the welfare of the community by applying the principles of social justice; and (4) Strengthening the community through the creation of social cohesion, inclusion, and optimizing the use of social capital that is built in the community. Social capital and social cohesion are developed to support the development of resilient cities through the interaction of social-ecological systems towards improving the quality of life of the community [87,88].

## Conclusion

The dynamics of the development of new urban areas contribute to changes in the characteristics of rural areas and transportation systems and the establishment of a central hierarchy for urban services, leading to an increase in land sales transactions carried out by community developers. Changes in the status and use value of agricultural land towards urban land use values for development contribute to an increase in investment flows towards changes in social structure, differences in status and economic strata and changes in people’s lifestyles. Changes in spatial characteristics have an impact on changes in land use forms, building characteristics, residential typology and changes in social interaction and social adaptation. These changes have triggered the socio-economic dynamics of society, economic disparities in wealth, segregation, and the emergence of exclusive societies. Land conversion coupled with housing and infrastructure development is positively associated with the integration of the Mamminasata Metropolitan urban system and is a determining factor causing environmental degradation.

The decline in environmental quality is the effect of changes in land use, increased socio-economic activities, infrastructure development, conversion of forest use and utilization of water catchment areas, which have an impact on changes in the flood cycle in suburban areas and the risk of natural disasters. Excessive use of groundwater has the potential to threaten environmental sustainability and balance. The complexity of the use of space and the availability of land for green open spaces that are converted to the needs of the development of socio-economic activities and housing development have a direct effect on the stability of the ecosystem. Efforts are needed to regulate the area of a residential building that functions to cover the ground surface, to ensure groundwater infiltration or the availability of ground water and to regulate the ground surface that is not covered by buildings to receive direct sunlight so that the soil can dry out and the air created around the building does not become polluted. moist. Thus, changes in land use, housing development, infrastructure development, and land reclamation simultaneously affect the environmental degradation of the new city area.

The spatial dynamics and integration of the Mamminasata Metropolitan urban system as part of the impact of the development of the new city area requires action to handle and control development towards the implementation of sustainable development. The strategic actions needed to deal with this are tightening the use of water catchment areas, setting up building boundaries, road borders, road-owned areas and road monitoring areas, limiting the development of horizontal housing and settlements to vertical development, integrating balanced housing and settlement development. through adequate infrastructure support, limiting development to productive land and forest areas through zoning arrangements and applying the principle of incentives and implementing the compact city concept by considering three important aspects, namely saving the environment, implementing the green economy concept, and involving community participation in monitoring environment. The sustainability of the city area has just been developed towards the implementation of the concept of environmental sustainability, namely the effectiveness and efficiency of land use and natural disaster mitigation. The direction of economic sustainability is oriented towards several principles, namely creating community economic competitiveness, preparing decent jobs, and encouraging strategic economic performance improvements through support for improving basic infrastructure performance, and minimizing economic and social disparities through creating equal opportunities for the community. The direction of social sustainability is oriented towards several principles, namely the creation of intergenerational equality and justice, the provision of optimal social services, ensuring the welfare of the community by applying the principles of social justice, and strengthening the community through the creation of social cohesion, inclusiveness, and optimizing the use of social capital that has been built in people’s lives.

## Supporting information

**S1 Table.** Spatial zoning of the new city area of Moncongloe–Pattalassang in 2021

(DOCX)

**S2 Table.** Space utilization of the new city area of Moncongloe–Pattalassang in 2021

(DOCX)

**S3 Table:** Changes in land use and development activities in the new city area of Moncongloe–Pattalassang

(DOCX)

## Acknowledgments

We are grateful for the participation of stakeholders in contributing ideas for the implementation of this study. We would like to thank the Ministry of Education, Culture, Research and Technology of the Republic of Indonesia and the University of Bosowa Foundation for their support and financial assistance in carrying out this research.

## Author Contributions

Conceptualization: Batara Surya.

Formal analysis: Batara Surya, Agus Salim, Syahrul Sariman.

Investigation: Batara Surya, Syahrul Sariman, Haeruddin Saleh.

Methodology: Batara Surya, Agus Salim, Seri Suriani, Hernita Hernita.

Supervision: Syahrul Sariman, Seri Suriani, Nasrullah Nasrullah, Haeruddin Saleh.

Writing – original draft: Batara Surya, Nasrullah Nasrullah, Hernita Hernita.

Writing – review & editing: Batara Surya, Nasrullah Nasrullah, Emil Salim Rasyidi

## Notes

### Competing Interest Statement

NO authors have competing interests Enter: The authors have declared that no competing interests exist.

